# Discovery and Biosynthesis of Sulfenicin and Its New-to-Nature Acylsulfenic Acid Functional Group

**DOI:** 10.1101/2024.10.16.618611

**Authors:** Dan Xue, Hongbin Zou, Wei Lv, Michael D. Madden, Xiaoying Lian, Mingming Xu, Conor Pulliam, Ethan A. Older, Lukuan Hou, Andrew Campbell, Tristan de Rond, Takayoshi Awakawa, Chunhua Yuan, Bradley S. Moore, Jie Li

## Abstract

Life’s organic molecules are built with diverse functional groups that enable biology by fine tuning intimate connections through time and space. As such, the discovery of new-to-nature functional groups can expand our understanding of the natural world and motivate new applications in biotechnology and biomedicine. Herein we report the genome-aided discovery of sulfenicin, a novel polyketide-nonribosomal peptide hybrid natural product from a marine *Streptomyces* bacterium bearing a unique acylsulfenic acid functionality. Through a series of heterologous biosynthesis, functional genetics, and enzymatic reconstitution experiments, we show that this previously described synthetic functional group is biologically assembled by a set of enzymes from both primary and secondary metabolism, including a novel flavin-dependent *S*-hydroxylase that hydroxylates a thiocarboxylic acid’s sulfur atom. While the sulfenicin biosynthetic gene cluster is presently without parallel in public databases, acylsulfenic acid-encoding enzymes are widely distributed in bacterial genomes, implying that this labile functional group may similarly have a broad distribution among specialized metabolites.

## Introduction

Bacterial natural products (NPs) play key roles in dictating how their producing organisms interact with their environment and other microbiota^1,2^. Such biologically active compounds were historically discovered via activity screening, but the proliferation of publicly available microbial sequencing data has paved the way for genome mining techniques to discover biosynthetic gene clusters (BGCs) which encode novel biosynthetic enzymes that produce unique NPs^3–5^. One such class attracting interest is sulfur-containing compounds, which exhibit tremendous structural diversity and a rich array of biological activities – including antibiotic, anticancer, and anti-inflammatory functions^6–9^, which make them especially attractive for medical or pharmaceutical applications. Despite their potential as drug candidates, the biosynthesis of sulfur-containing NPs is understudied and lags behind the well-documented sulfur incorporation and trafficking mechanisms in primary metabolism. Different mechanisms (e.g., substitution reactions with sulfur nucleophiles^10–13^, additions^14^, and radical reactions^15^) catalyzed by a variety of enzymes (e.g., *S*-transferases^16,17^, thioesterases^18^, cyclases^19,20^, oxygenases^21–23^, and radical *S*-adenosylmethionine enzymes^24^) have been recently identified from only a small fraction of sulfur-containing NPs^25^, thus leaving much of their biosynthesis unexplored.

Here we report the (i) identification of a unique BGC in a marine bacterium *Streptomyces* sp. CNT360, for which the final products could not be predicted; (ii) subsequent discovery of its resulting NP sulfenicin, which contains a new-to-nature acylsulfenic acid functional group; (iii) *in vivo* and *in vitro* elucidations of the acylsulfenic acid’s biosynthesis involving seven proteins/enzymes from both primary and secondary metabolism; and (iv) characterization of an unprecedented flavin-dependent *S*-hydroxylase in acylsulfenic acid biosynthesis. Furthermore, we note that the three clustered genes encoding an acyl-CoA synthetase, a CoA-transferase, and a novel *S*-hydroxylase may represent an acylsulfenic acid gene cassette that is widely distributed in bacterial genomes, implying that the newly described acylsulfenic acid functional group may be readily introduced into various core structures of other NPs.

## Results and discussion

### Discovery of a new-to-nature acylsulfenic acid functional group

We performed microbial genome mining to discover new NPs biosynthetic pathways involving sulfur-related biosynthetic genes, especially those clustered with genes encoding novel tailoring enzymes. Our effort identified a BGC, which we named *taa* after its subsequently identified thiazole <u>a</u>cylsulfenic <u>a</u>cid product, in a marine bacterium, *Streptomyces* sp. CNT360. The *taa* BGC encodes a polyketide synthase-nonribosomal peptide synthetase (PKS-NRPS) hybrid and several putative tailoring enzymes, two of which had notable homology to previously reported enzymes involved in thiocarboxylic acid biosynthesis^26^ (Fig. 1a and Supplementary Table S1). Searches in all publicly available genome databases revealed no other similar BGCs, suggesting that *taa* was a singleton cluster likely encoding novel sulfur containing compounds. To probe for any potentially unknown products of this BGC, we captured the *taa* BGC by transformation-associated recombination (TAR) cloning^27,28^ and integrated this BGC into the genome of the heterologous expression strain *S. coelicolor* M1152, generating M1152_*taa* that harbors the *taa* BGC (Supplementary Tables S2 and S3). Liquid chromatography photodiode array detection mass spectrometry (LC-PDA-MS) analysis of the ethyl acetate extract of M1152_*taa* revealed two major products, named carboxy-presulfenicin (**1**) and sulfenicin (**2**) (Fig. 1b), which exhibited distinct UV absorbance profiles (Fig. 1c). Upon isolation, the structure of carboxy-presulfenicin (**1**) was elucidated (Supplementary Notes), using a combination of high-resolution electrospray ionization mass spectrometry (HR-ESIMS) (Supplementary Fig. S1), infrared spectroscopy (IR) (Fig. 1c), and nuclear magnetic resonance spectroscopy (NMR) (Supplementary Figs. S2-S6 and Table S4). The structure of carboxy-presulfenicin **(1**) was further confirmed by X-ray diffraction analysis of its complex with Cu(II), which also established the stereochemistry of the polypropionate acyl side chain in **1** (Fig. 1d and Supplementary Table S5).

**Fig 1.**
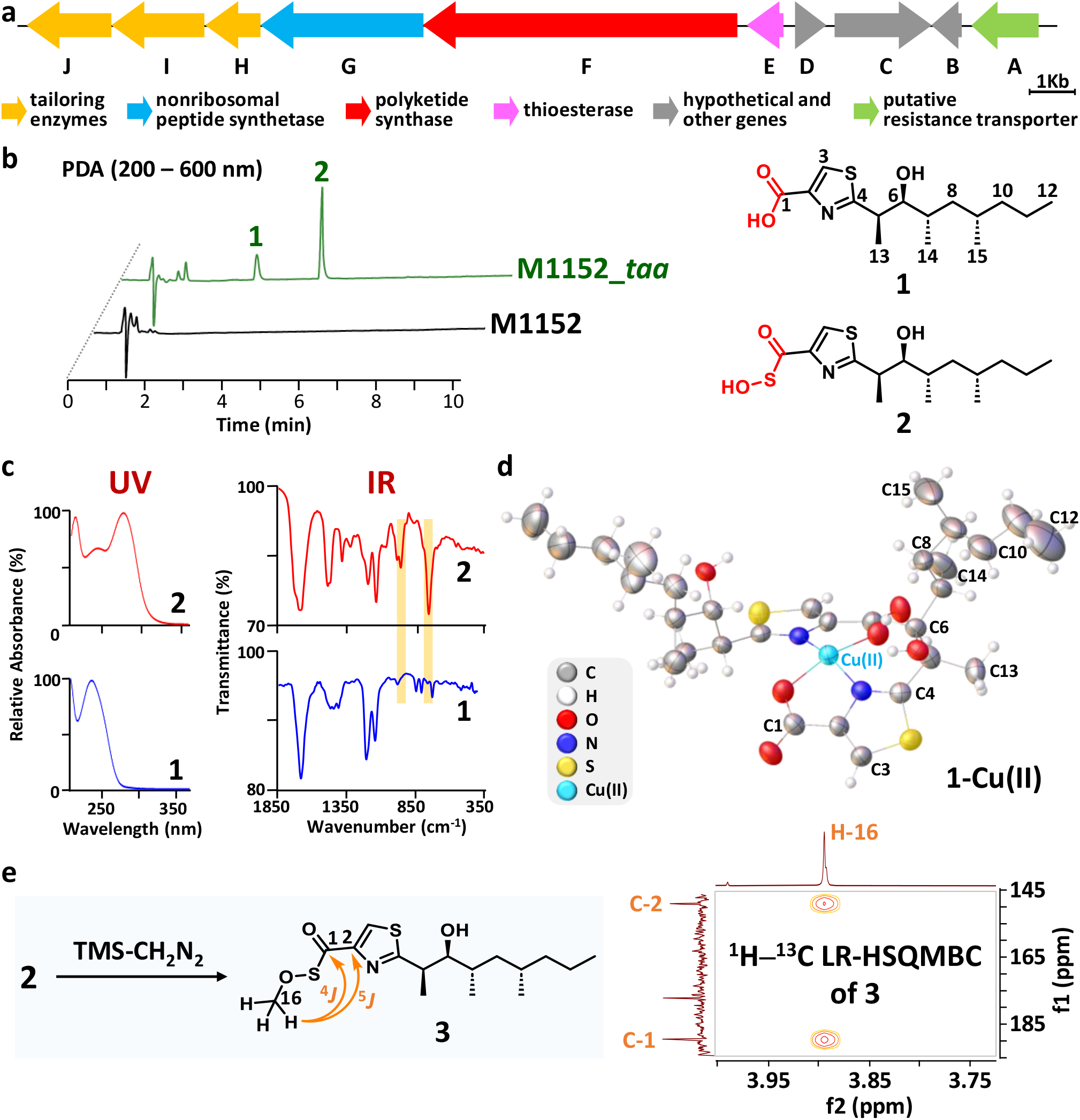
Heterologous expression of the *taa* BGC and structural characterization of a novel thiazole acylsulfenic acid. **a**, The *taa* BGC in *Streptomyces* sp. CNT360 encodes a hybrid PKS-NRPS (TaaE, TaaF, and TaaG) and three putative tailoring enzymes (TaaH: Group B flavoprotein monooxygenase (FPMO); TaaI: CoA-transferase; TaaJ: Acyl-CoA synthetase) responsible for the biosynthesis of a novel thiazole acylsulfenic acid. **b**, LC-PDA analysis of ethyl acetate extract of the supernatant from the heterologous expression strain M1152_*taa* indicates the production of two new compounds, carboxy-presulfenicin (**1**) and sulfenicin (**2**), that are not produced by the wild-type M1152. **c**, UV absorbance profiles and IR spectra of carboxy-presulfenicin (**1**) and sulfenicin (**2**). The UV profiles indicated *λ*_max_ shift from 236 nm in carboxy-presulfenicin (**1**) to 282 nm in sulfenicin (**2**). The IR spectra indicated a C-S vibration at 971.9 cm^-1^ and a S-O vibration at 779.1 cm^-1^ for sulfenicin (**2**), both of which are absent in the IR spectrum of carboxy-presulfenicin (**1**). **d**, X-ray crystal structure of two molecules of carboxy-presulfenicin (**1**) coordinated with Cu(II). **e**, Methylation of sulfenicin (**2**) by TMS-CH_2_N_2_ yielded **3**, allowing for the observation of key ^4^*J* and ^5^*J* correlations in the ^1^H–^13^C LR-HSQMBC NMR spectrum recorded in CDCl_3_.

Remarkably, HR-ESIMS analysis of sulfenicin (**2**) generated an [M + H]^+^ ion with a *m*/*z* 332.1342 (Supplementary Fig. S1), suggesting a molecular formula of C_15_H_25_NO_3_S_2_ that notably contained an additional sulfur atom in comparison with carboxy-presulfenicin (**1**) (C_15_H_25_NO_3_S). In the ^13^C NMR spectrum of isolated sulfenicin (**2**), the C=O chemical shift was observed at *δ*_C_ 191.8 ppm (Supplementary Figs. S7-S13 and Supplementary Table S6), which was much further downfield than the carboxyl C=O chemical shift of carboxy-presulfenicin (**1**) (*δ*_C_ 164.2 ppm; Supplementary Fig. S3). This observation indicated that the additional sulfur atom is directly bonded to the C=O group, inducing a deshielding effect from the partial positive charge present on the sulfur (RC(O^-^)=S^+^R’). Furthermore, compared to carboxy-presulfenicin (**1**), sulfenicin (**2**) displayed an extra IR band at 971.9 cm^-1^, attributed to the C-S vibration, and an additional characteristic S-O vibration band at 779.1 cm^-1^ (Fig. 1c)^29,30^. Collectively, these data suggested that sulfenicin (**2**) contains an acylsulfenic acid group in place of the carboxylic acid group in carboxy-presulfenicin (**1**). To provide further support, we treated sulfenicin (**2**) with a solution of TMS-CH_2_N_2_ in CH_2_Cl_2_ to generate compound **3**, forming a terminal methyl group (Fig. 1e, Supplementary Figs. S14-S19, Supplementary Table S7, and Supplementary Notes). Indicated by the ^1^H–^13^C long-range heteronuclear single quantum multiple bond correlation (LR-HSQMBC), we measured key correlations between newly added methyl H-16 (*δ*_H_ 3.89 ppm) to C-1 (*δ*_C_ 189.4 ppm) of the carbonyl group and to C-2 (*δ*_C_ 149.2 ppm), thereby confirming the presence of the acylsulfenic acid functional group.

A pyridine-bearing natural product previously detected in *Pseudomonas* spp. was hypothesized to contain an acylsulfenic acid group based on chemical analysis, but was never isolated^31^. Thus, chemical derivatizations were used to methylate the acylsulfenic acid group or oxidize a thiocarboxyl group to an acylsulfenic acid group for characterization^31,32^. The present study represents the first direct isolation and characterization of a natural product containing the unique acylsulfenic acid functional group. Interestingly, while the isolated sulfenicin (**2**) was not stable in its purified form, it remained stable and was produced in high yield during cultivation of both the native producer *Streptomyces* sp. CNT360 and the heterologous expression strain M1152_*taa*, suggesting that certain components present during the cultivation could stabilize the acylsulfenic acid product. As such, this stable form may exert biological functions in its producing organism.

### *In vivo* biosynthetic pathway interrogation of sulfenicin

To investigate the biosynthesis of the unique acylsulfenic acid-containing sulfenicin (**2**), we first used a gene deletion strategy to probe the functions of the critical enzymes encoded by the *taa* BGC. The heterologous expression strain with *taaA-D* deleted still produced carboxy-presulfenicin (**1**) and sulfenicin (**2**). Thus, we concluded that *taaA-D* are not essential biosynthetic genes (Supplementary Figs. S20 and S21). We hypothesized that sulfenicin (**2**) is biosynthesized from carboxy-presulfenicin (**1)**, while carboxy-presulfenicin (**1**) was envisioned to be synthesized by a canonical modular PKS (TaaF)-NRPS (TaaG) hybrid enzymatic assembly line and offloaded by a thioesterase (TE, TaaE) (Fig. 2a). Accordingly, after deleting *taaE*, the production of carboxy-presulfenicin (**1**) and sulfenicin (**2**) in the heterologous expression strain was completely abolished (Fig. 2b). Thus, we focused our analysis on the three remaining downstream biosynthetic genes in the BGC – *taaH*-*J*. Deletion of *taaI* or *taaJ* each abolished the production of sulfenicin (**2**) while increasing the production of carboxy-presulfenicin (**1**), which suggested that TaaJ and TaaI are involved in the conversion of carboxy-presulfenicin (**1**) to sulfenicin (**2**). Deletion of *taaH* also abolished the production of sulfenicin (**2**) with concomitant generation of two new compounds, **4** and **5** (Fig. 2b). Genetic complementation of *taaH* into the M1152_*taa*Δ*taaH* deletion mutant restored the production of sulfenicin (**2**) while eliminating **4** and **5** (Fig. 2b), further confirming TaaH’s role in generating sulfenicin (**2**). Based on these results, we speculated that **4** and/or **5** were possible additional substrates of TaaH for conversion to sulfenicin (**2**). We isolated the more abundant **4** and elucidated its chemical structure by NMR (Supplementary Figs. S22-S26, Supplementary Table S8, and Supplementary Notes). Since compounds **4** and **5** shared the same MS2 fragmentation pattern but only differed by 14 Da in their exact masses (Supplementary Fig. S27), we reasoned that **5** could be the *S*-demethylated version of **4**. We treated **4** with potassium hydrosulfide to generate a corresponding thiocarboxylic acid product (Fig. 2c) and confirmed its structure by NMR (Supplementary Figs. S28-S31, Supplementary Table S9, and Supplementary Notes). This resulting product showed identical retention time, exact mass, and MS2 fragmentation patterns to **5** (Supplementary Fig. S27). Thus, we identified **5** as the thiocarboxylic acid version of **4**. The structural resemblance and possible biosynthetic conversion between the thiocarboxylic acid and acylsulfenic acid led to our hypothesis that **5**, named thio-presulfenicin, was the true intermediate used to generate sulfenicin (**2**), while methylation of thio-presulfenicin (**5**) to produce **4** – either enzymatic or non-enzymatic – could be a possible detoxification strategy as high concentrations of thiocarboxylic acid or thiol-containing compounds may be toxic^33^.

**Fig 2.**
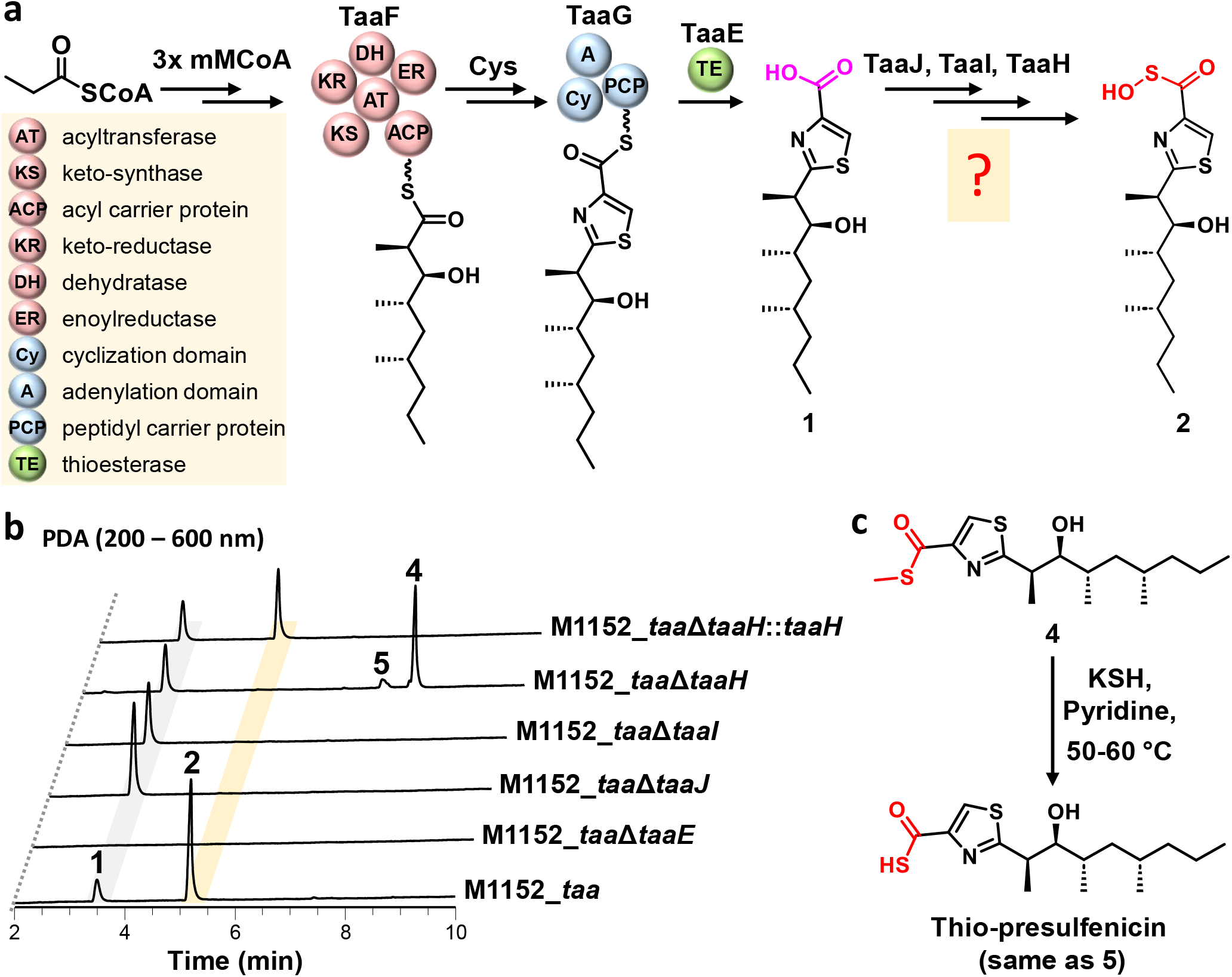
*In vivo* deletion of genes in M1152_*taa* to probe biosynthetic function. **a**, The hypothesized biosynthetic pathway of sulfenicin (**2**). TaaF, a putative iterative type I PKS, is expected to initiate assembly with propionyl-CoA and then elongate three times with methyl malonyl-CoA, reducing and dehydrating each module as necessary to form the aliphatic chain seen in carboxy-presulfenicin (**1**). TaaG, a putative NRPS, is then proposed to incorporate and subsequently cyclize L-cysteine to form the thiazole moiety in carboxy-presulfenicin (**1**). Once the backbone is assembled, TaaJ, TaaI, and TaaH are predicted to incorporate a sulfur and hydroxylate it to form sulfenicin (**2**). **b**, PDA chromatograms of sequential *in vivo* gene deletion of *taa* BGC, suggesting that TaaI and TaaJ are involved in formation of the thiocarboxylic acid-containing intermediate thio-presulfenicin (**5**), while TaaH is predicted to introduce the novel acylsulfenic acid moiety and form sulfenicin (**2**). Carboxy-presulfenicin (**1**) and sulfenicin (**2**) in the chromatograms were identified based on retention time, exact mass, and MS2 fragmentation compared to standards (Supplementary Figs. S1 and S21). Each PDA chromatogram is normalized to the same scale. **c**, Chemical synthetic route to generate thio-presulfenicin (**5**) from compound **4**.

Taken together, our *in vivo* gene deletion results indicated that the acyl-thiazole skeleton of carboxy-presulfenicin (**1**) and sulfenicin (**2**) was produced by a PKS-NRPS hybrid assembly line (TaaE-G), while TaaH-J are involved in converting the carboxylic acid to form the novel acylsulfenic acid functional group in sulfenicin (**2**). This biosynthetic process involves a cryptic thiocarboxylic acid intermediate (thio-presulfenicin, **5**) that was not originally detected from the native or heterologous expression strain, but only from the M1152_*taa*Δ*taaH* deletion mutant, suggesting that the intermediate thio-presulfenicin (**5**) is robustly converted to the final product sulfenicin (**2**) during biosynthesis.

### *In vitro* enzymatic reconstitution of the acylsulfenic acid functional group involves seven proteins/enzymes from both primary and secondary metabolism

We next investigated the individual functions of TaaH, TaaI, and TaaJ. Based on the *in vivo* gene deletion results and enzyme annotations, we hypothesized that TaaJ (Acyl-CoA synthetase) first activates the carboxylic acid in carboxy-presulfenicin (**1**) using CoA in an ATP-dependent manner, forming **1-CoA** (Fig. 3a). Afterwards TaaI (CoA-transferase) transfers a sulfur atom to **1-CoA** to form the thiocarboxylic acid in thio-presulfenicin (**5**), analogous to the biosynthesis of thioplatensimycin (thioPTM) (Fig. 3a)^26^. Finally, an enzymatic *S*-hydroxylation of thio-presulfenicin (**5**) by TaaH, a group B flavoprotein monooxygenase (FPMO), generates the acylsulfenic acid functional group in sulfenicin (**2**) (Fig. 3a).

**Fig 3.**
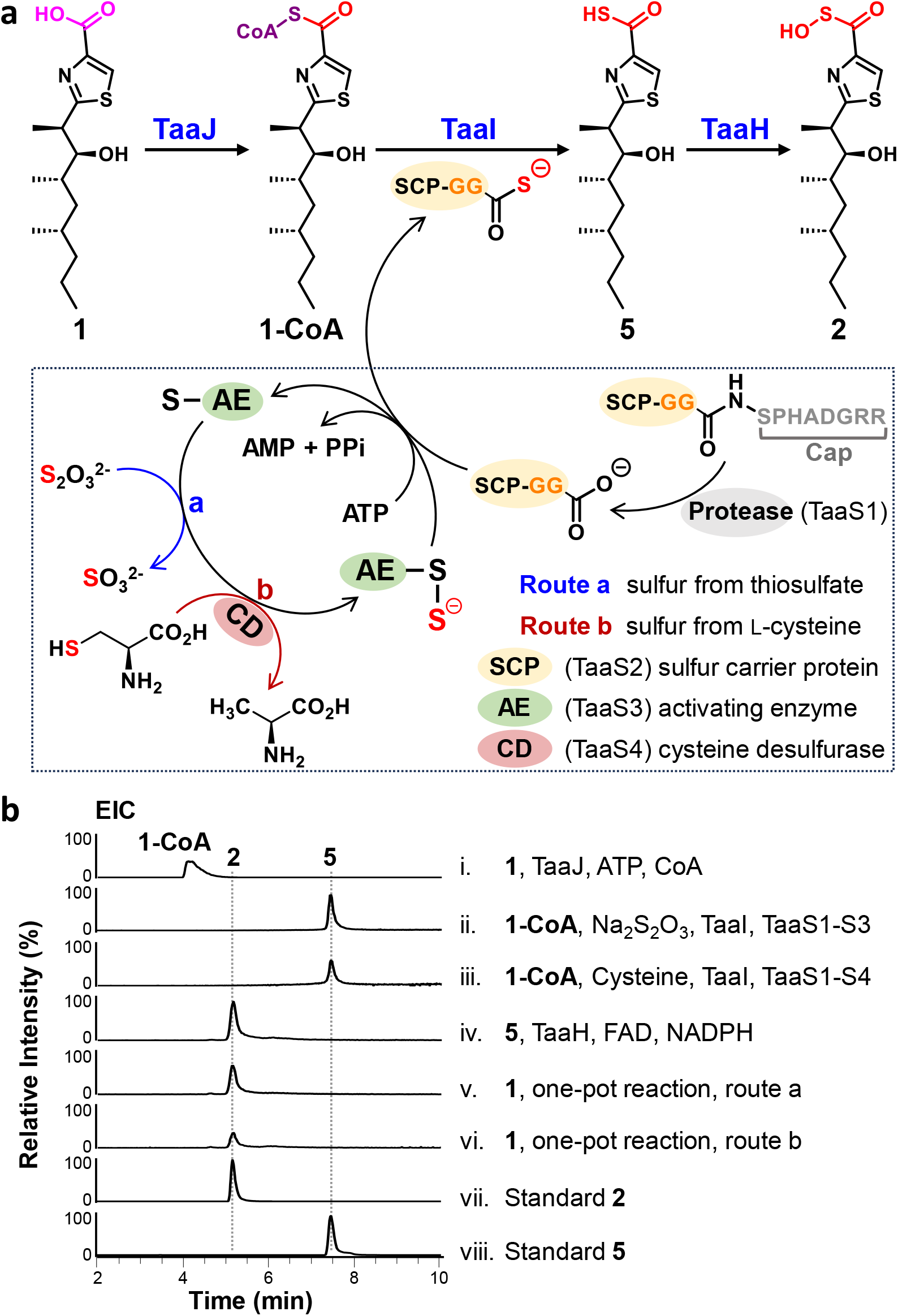
*In vitro* reconstitution of acylsulfenic acid-synthesizing enzymes. **a**, Hypothesized biosynthetic pathway for the acylsulfenic acid functional group in sulfenicin (**2**). TaaJ first generates **1-CoA** from carboxy-presulfenicin (**1**), after which TaaI incorporates sulfur from primary metabolic pathways to yield thio-presulfenicin (**5**), which is finally hydroxylated by TaaH to yield the final product, sulfenicin (**2**). **b**, *In vitro* reconstitution of sulfenicin (**2**)’s biosynthesis. Extracted ion chromatograms (EICs) for **1-CoA** (*m*/*z* 1049.2670, [M + H]^+^), thio-presulfenicin (**5**) (*m*/*z* 316.1394, [M + H]^+^), and sulfenicin (**2**) (*m*/*z* 332.1342, [M + H]^+^) are depicted. Sulfur from sodium thiosulfate (Traice v, route a) and L-cysteine (Trace vi, route b), arising from primary metabolism, is incorporated into the final acylsulfenic acid.

To explore this hypothesis, we first considered the source of sulfur. The lack of any bioinformatically-predicted sulfur-donating genes in the *taa* BGC led us to speculate that the sulfur atom is incorporated into **1-CoA** from primary metabolism^34^ to form the thiocarboxylic acid in thio-presulfenicin (**5**). We considered two potential routes of sulfur incorporation from primary metabolism: from inorganic and organic sources (Fig. 3a). Our feeding of ^34^S-labeled L-cysteine indeed led to the incorporation of two ^34^S into the final product sulfenicin (**2**) (Supplementary Fig. S32), supporting the incorporation of sulfur from primary metabolism.

Next, we enzymatically reconstituted sulfenicin (**2**) *in vitro* in a systematic manner. We first incubated TaaJ with carboxy-presulfenicin (**1**), ATP and CoA as substrates as well as MgCl_2_ as a cofactor. We detected the production of **1-CoA** as well as its corresponding phosphopantetheine (PPant) ejection ion (Trace i in Fig. 3b and Supplementary Fig. S33 and S34). Next, we used **1-CoA** as a substrate to establish the enzymatic activity of TaaI. As a transferase, TaaI requires another substrate to donate its sulfur to **1-CoA** and form the thiocarboxylic acid product. Via a homology search, we identified one JAMM metalloprotease (TaaS1), four sulfur carrier proteins (SCPs), one activating enzyme (TaaS3), and one cysteine desulfurase (TaaS4) in *Streptomyces* sp. CNT360’s genome based on their similarities to previously reported proteins and enzymes used for sulfur delivery from primary metabolism (Supplementary Table S10)^26,34,35^. We focused on a particular SCP (TaaS2) as it had the highest sequence similarity to an identified SCP^26^ and its encoding gene was clustered with TaaS1’s gene. TaaS2 had capped C-termini, and SCPs with such feature typically have higher activities than those with pre-exposed C-termini^34^, but they require a protease to remove the cap and activate the enzyme (Fig. 3a)^35^. As such, TaaS2’s clustering with a protease suggested that it had high potential to be an active SCP. After generating **1-CoA** *in situ* and adding sodium thiosulfate (Na_2_S_2_O_3_) as an inorganic sulfur-donating substrate, we incubated the mixture with TaaI and TaaS1-S3 to generate thio-presulfenicin (**5**) (Traces ii and viii in Fig. 2c and Supplementary Fig. S27). Furthermore, we performed a similar assay, replacing Na_2_S_2_O_3_ with L-cysteine as an organic sulfur donor and additionally adding TaaS4 to cleave the sulfur from L-cysteine. Likewise, we observed the formation of thio-presulfenicin (**5**) (Trace iii in Fig. 3b and Supplementary Fig. S27).

We next sought to establish the enzymatic activity of TaaH using thio-presulfenicin (**5**) as a substrate. Since our bioinformatic analysis annotated TaaH as a group B FPMO, we first confirmed its flavoprotein identity by evaluating the binding of FAD and NADPH to TaaH. UV-vis spectroscopy of TaaH showed prominent absorbances at 380 nm and 450 nm, indicating the presence of bound FAD in the enzyme; the NADPH binding to TaaH was confirmed by adding free NADPH and observing the absorbance at 340 nm (Supplementary Fig. S35). Next, we incubated TaaH with thio-presulfenicin (**5**) as a substrate along with FAD and NADPH as cofactors to generate the final acylsulfenic acid product, sulfenicin (**2**) (Traces iv and vii in Fig. 3b and Supplementary Fig. S21). Using NADH as an alternative to NADPH also yielded sulfenicin (**2**), but at approximately 50% lower yield (Supplementary Fig. S36). We also observed that thio-presulfenicin (**5**) could be slightly non-enzymatically oxidized into sulfenicin (**2**) under a prolonged incubation. However, this non-enzymatic process was ∼125 times slower than the oxidation by TaaH, supporting our identification of TaaH’s enzymatic activity. Finally, we conducted two sets of one-pot *in vitro* enzymatic syntheses of sulfenicin (**2**) from carboxy-presulfenicin (**1**) using either Na_2_S_2_O_3_ or L-cysteine as the sulfur donor. In the first one-pot reconstitution, we combined six proteins/enzymes (TaaH-J and TaaS1-S3) with the substrates carboxy-presulfenicin (**1**), ATP, CoA and Na_2_S_2_O_3_, along with cofactors FAD, NADPH, and MgCl_2_ (Route a in Fig. 3a) to successfully produce sulfenicin (**2**) (Trace v in Fig. 3b). Similarly, in the second one-pot reconstitution, L-cysteine was used as the sulfur donor instead of Na_2_S_2_O_3_ (Route b in Fig. 3a) and seven proteins/enzymes (TaaH-J and TaaS1-S4) were involved to generate sulfenicin (**2**) (Trace vi in Fig. 3b). Taken together, our *in vitro* enzymatic reconstitutions confirm the multi-enzymatic process of converting a carboxyl group to an acylsulfenic acid group. The energy investment and coordination requirement for incorporation of seven proteins/enzymes from both primary and secondary metabolism to generate an acylsulfenic acid functional group suggests a potentially important biological role of this functional group in nature.

### Analysis and characterization of TaaH, an unprecedented flavin-dependent *S*-hydroxylase

After establishing the biosynthetic pathway of sulfenicin (**2**), we specifically characterized TaaH’s enzymology due to its *S*-hydroxylation activity. We carried out a steady-state kinetic analysis of TaaH using thio-presulfenicin (**5**) as the substrate, which revealed a catalytic constant (Kcat) of 29.8 min^-1^, a maximum reaction velocity (Vmax) of 0.012 μM/s, and a Michaelis constant (Km) of 46.6 μM (Fig. 4a). The Km value demonstrates moderate substrate affinity and the Kcat shows appropriate enzymatic turnover, supporting that the hydroxylation of thio-presulfenicin (**5**) by TaaH is an enzymatic process. Among the three families of group B FPMOs, we first compared TaaH’s sequence with known flavin-containing monooxygenases (FMOs) from various kingdoms to identify TaaH’s potential active residues. Sequence alignment with other known FMO proteins showed that TaaH contains previously reported NADPH and FAD binding sites (GxGxxG) but does not feature the conserved FMO motif (FxGxxxHxxx(Y/F)) (Supplementary Fig. S37 and Table S11)^36^. This region in TaaH is xxxxxxHxxxY, which lacks the FxG compared to the typical FMO motif. Despite this difference, TaaH still resembles FMOs more closely than the two other families of group B FPMOs, namely Baeyer-Viliger monooxygenases (BVMOs) and *N*-hydroxylating monooxygenases (NMOs). In BVMOs, the C-terminal amino acid of this region is tryptophan (W) while NMOs are characterized by the presence of a single conserved histidine residue (xxxxxxHxxxx), neither of which are featured in TaaH (Supplementary Fig. S37 and Table S11). Furthermore, previously reported active site residues of FMOs – such as N/H62, E/D176, and R202 – are not conserved in TaaH (Supplementary Fig. S37)^37^. Due to the absence of the conserved FMO motif and active site residues, we examined the structure of TaaH through molecular modeling. A homology model of TaaH was constructed based on the crystal structures of monooxygenases. Subsequently, the interaction between thio-presulfenicin (**5**) and TaaH was investigated using molecular docking. The docking results revealed that thio-presulfenicin (**5**) binds to the active site of TaaH, in proximity to both cofactors NADPH and FAD (Fig. 4b). Examination of the residues making up the binding pocket showed two typical Rossmann fold motifs (GxGxxG) situated proximal to the FAD and NADPH (Fig. 4c)^37,38^. The H150 and Y154 residues (black) – part of the FMO motif (FxGxxxHxxx(Y/F)) – are situated within the binding pocket but are far from the substrate-cofactor complex (Fig. 4c). Finally, F53, P131, P320, W321, and Q322 (red) are arranged closest to the substrate and not conserved across other selected FMOs (Fig. 4c and Supplementary Fig. S37).

**Fig 4.**
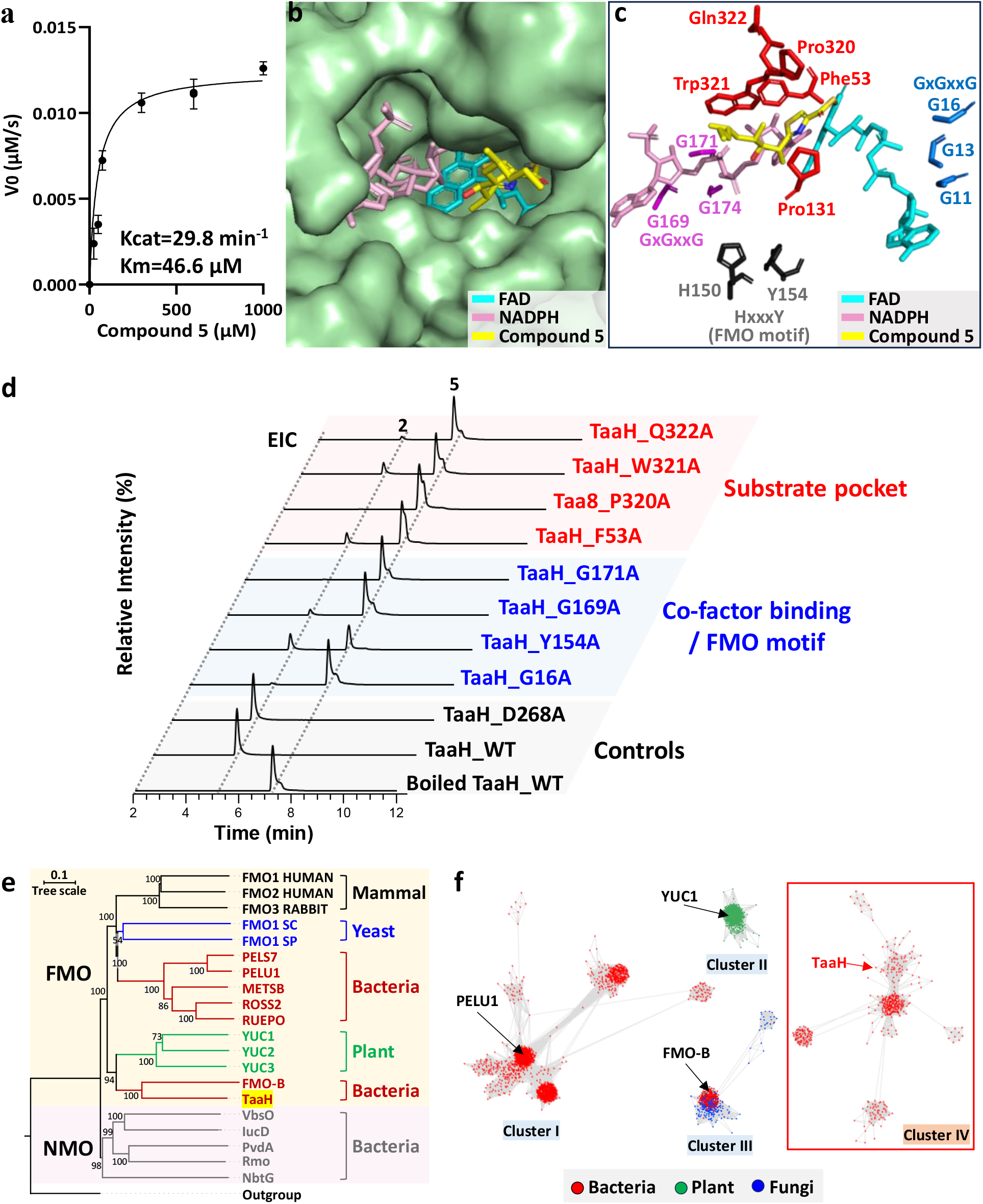
Analysis and characterization of TaaH. **a**, Steady-state kinetic analysis of TaaH with thio-presulfenicin (**5**) as a substrate. **b**, Molecular docking using thio-presulfenicin (**5**) as a substrate (yellow) as well as NADPH (pink) and FAD (cyan) as cofactors in TaaH’s binding pocket. **c**, Molecular docking of TaaH with its substrate, thio-presulfenicin (**5**) (yellow), and cofactors, NADPH (pink) and FAD (cyan). Residues involved in cofactor stabilization (magenta and blue), FMO motif (black), and substrate interaction (red) are shown. **d**, *In vitro* biosynthetic analyses using TaaH mutants. Peaks indicating the presence of sulfenicin (**2**, *m/z* 332.1342, [M + H]^+^) and thio-presulfenicin (**5**, *m/z* 316.1394, [M + H]^+^) are depicted on the EICs. Sulfenicin (**2**) and thio-presulfenicin (**5**) in the chromatograms were identified based on retention time, exact mass, and MS2 fragmentation. **e**, Phylogenetic analysis of TaaH places it among known *S*-oxidizing enzymes (FMO-B) and *N*-hydroxylating enzymes (plant YUCCA proteins and bacterial NMOs) while distancing it from type I bacterial FMOs. **f**, Sequence similarity networking (SSN) of TaaH with the UniProt database clusters the enzyme in a group (Cluster IV, ∼500 nodes) distinct from type I FMOs (Cluster I, ∼2,500 nodes), plant FMOs/YUCCA proteins (Cluster II, ∼1,100 nodes), and type II FMOs (Cluster III, ∼700 nodes). Cluster IV also includes uncharacterized enzymes from diverse phyla.

To probe the function of the residues identified in the molecular docking analysis, we individually mutated each of these residues to alanine and subjected these mutants to *in vitro* enzymatic assays (Supplementary Fig. S38). Mutating G16, G169, or G171 severely decreased or abolished the production of sulfenicin (**2**) (Fig. 4d), suggesting that they play important roles in binding the FAD and NADPH cofactors to the pocket. Similarly, mutating the residues of the FMO motif either prevented the expression of soluble TaaH (H150A) or diminished sulfenicin (**2**)’s production (Y154A) (Fig. 4d). This region is considered to link the FAD and NADPH binding pockets of FMOs, ensuring correct domain rotations and conformational changes^39^. Mutations in this region have also been reported to decrease the solubility of the enzyme or inactivate it^36,40^. Next, we mutated each of the unique residues closest to the substrate (F53, P320, W321, Q322, and P131) to alanine individually. According to the docking results, all of these residues are positioned within 5.0 Å of thio-presulfenicin (**5**). They may directly interact with the substrate, or NADPH, FAD cofactors, as exemplified by the potential π-π stacking between F53 and the thiazole ring of thio-presulfenicin (**5**), as well as the hydrophobic interactions involving P320, W321, Q322, and P131 with the aliphatic chain of thio-presulfenicin (**5**). The individual TaaH mutant of F53A, P320A, W321A, and Q322A diminished the production of sulfenicin (**2**), suggesting that they each play important roles in catalysis (Fig. 4d). The P131A TaaH mutant failed to express, suggesting that this residue is required to achieve the correct protein architecture (Fig. 4c). To avoid the possibility that any decreased activities observed arose from improper protein folding, we measured the circular dichroism (CD) spectra of the mutants in comparison with the wild-type (WT) TaaH to ensure the correct protein folding before testing the enzymatic activity of these mutants (Supplementary Fig. S38). We also randomly mutated an amino acid outside of the catalytic pocket (D268A) as an additional comparison. This mutant showed similar activity to the WT TaaH in the *in vitro* assay (Fig. 4d), further supporting that the modified enzymatic activities of the mutants described above were derived from the roles of those residues in TaaH’s catalysis.

TaaH’s unprecedented ability to hydroxylate a thiocarboxylic acid moiety also prompted us to consider how the enzyme acquired this activity during evolution. Thus, we performed phylogenetic analysis of TaaH with other group B FPMOs (Fig. 4e and Supplementary Table S11). TaaH is grouped with FMO-B, a type II FMO (Fig. 4e). This subclass is known to catalyze the oxidation of various *S*-containing compounds, such as thioanisole^41,42^. However, the hydroxylation of sulfur has not yet been reported for this subclass. Type II FMOs are also considered to be an ancestor of a plant FMO subclass, the YUCCA proteins (YUCs)^43^. The close phylogeny of TaaH to the plant YUCs (Fig. 4e) is notable, as YUCCA proteins catalyze *N*-hydroxylation reactions^44^, similar to the *S*-hydroxylation seen in TaaH. The only other family within the group B FPMOs superfamily that also catalyzes *N*-hydroxylations are NMOs, which are usually found in microorganisms^36^. In agreement with these reported activities, the phylogenetic tree indicates that TaaH is located close to both YUCCA proteins and bacterial NMOs while comparatively distant from the type I FMOs found in bacteria, as well as FMOs from yeast and mammals (Fig. 4e). Interestingly, TaaH’s placement forms a unique clade capable of the heteroatom-hydroxylation retained from the YUCCA proteins and NMOs, but instead using sulfur as a substrate like FMO-B. We then used sequence similarity networking (SSN) to examine the prevalence of TaaH homologs across nature. The top 9,703 sequences blasted from the UniProt database with an *e*-value threshold of 10^−10^ and an alignment score of 91 were clustered and evaluated (Supplementary Fig. S39), a subset of which is pictured in Fig. 4f. In the SSN, TaaH is not grouped with any known bacterial type I FMOs (Cluster I), plant FMOs/YUCCA proteins (Cluster II), or type II FMOs (Cluster III), supporting its status as a representative member of a new group of FMOs. TaaH’s cluster (Cluster IV) includes sequences from diverse phyla such as Actinobacteria, Firmicutes, and Bacteroidetes, indicating that TaaH homologs and enzymatic *S*-hydroxylation may exist broadly in nature. In particular, the four most closely related nodes to TaaH in Cluster IV each belongs to an acylsulfenic acid cassette in different BGCs (Supplementary Fig. S40), like *taaH-J* in *Streptomyces* sp. CNT360, suggesting that these BGCs may also encode an acylsulfenic acid functional group installed on other diverse core structures of natural products. We anticipate that the acylsulfenic acid functional group may chelate metals and convey metal-sequestering and/or antibiotic potential, a direction we are investigating.

## Conclusion

Sulfur-containing compounds have emerged as a promising class of NPs due to their structural diversity and potential medical applications. In this study, we discovered a novel acylsulfenic acid functional group in the sulfenicin family of natural products produced by an orphan BGC of a marine *Streptomyces* bacterium. The biosynthesis of this functional group involves seven different proteins/enzymes from both primary and secondary metabolism, including a flavin-dependent *S*-hydroxylase that is distinct from known flavoproteins^45^. Bioinformatic analysis also suggests that acylsulfenic acid-encoding enzymes might be more prevalent in bacteria than previously recognized, which could lead to the integration of acylsulfenic acid functional group into various NP core structures. This study enriches the understanding of sulfur-containing NP chemical diversity and biosynthesis^46^.

## Methods

### General experimental procedures

UV-vis spectra were obtained on a Biospectrometer^®^ basic UV-vis spectrophotometer (Eppendorf, Hamburg, DE). Infrared spectra (IR) were taken on a PerkinElmer spectrum 100 fourier transform infrared spectrometer incorporated with a diamond crystal plate as the reflector. Nuclear magnetic resonance (NMR) spectra were acquired on a Bruker Avance III HD 400MHz spectrometer with a 5 mm BBO 1H/19F-BB-Z-Gradient prodigy cryoprobe, a Bruker Avance III HD 600 MHz spectrometer with a PA BBO 500S2 BBF-H-D_05 Z SP probe, and a Bruker Avance III HD Ascend 850 MHz spectrometer equipped with a 5 mm triple-resonance Observe (TXO) cryoprobe with Z-gradients, controlled by TopSpin 3.6.1 software. Unless otherwise specified, all spectra were acquired at 25°C in solvents as specified in the text, with reference to residual ^1^H or ^13^C signals in the deuterated solvents. High-resolution electrospray ionization mass spectrometry (HR-ESIMS) spectra were obtained on a Thermo Scientific Orbitrap Velos Pro hybrid ion trap-orbitrap mass spectrometer by direct injection. Liquid chromatography photodiode array detection electrospray ionization mass spectrometry (LC-PDA-ESIMS) data were acquired on a Thermo Dionex Ultimate 3000 UHPLC system equipped with a diode array multiple wavelength detector and an LTQ XL linear ion trap mass spectrometer controlled by Thermo Xcalibur (version 4.2.47). The ion trap mass spectrometer was set at a sheath gas flow rate of 35 arbitrary units, a source heater temperature of 325°C, and a capillary temperature of 350°C. Semi-preparative HPLC purifications were performed using a Thermo Dionex Ultimate 3000 HPLC system with corresponding pump, autosampler, UV-vis detectors, fraction collectors, and Chromeleon software (version 7.2.10). LCMS grade and HPLC-grade CH_3_CN, H_2_O and formic acid were purchased from Fisher Scientific. Deuterated solvents were purchased from Cambridge Isotopes.

### General materials, strains, and culture conditions

Biochemicals and media components for bacterial cultures were purchased from Thermo Fisher Scientific Co. Ltd. (USA) unless otherwise stated. Restriction endonucleases were purchased from New England Biolabs, Inc. (USA). Chemical reagents were purchased from standard commercial sources. PCR amplifications were carried out on an Eppendorf^®^ Mastercycler^®^ Nexus X2 Thermal Cycler (Eppendorf Co., Ltd. Germany) using PrimeSTAR HS DNA polymerase (Takara Biotechnology Co., Ltd. Japan). The E.Z.N.A.^®^ Gel Extraction Kit (Omega Bio-tek, Inc., USA) was used for PCR products purification. The NEBuilder^®^ HiFi DNA Assembly master mix (New England Biolabs, Inc., USA) was applied for Gibson assembly. Oligonucleotide synthesis and DNA sequencing were performed by Eton Bioscience, Inc. (USA). All strains used in this study are listed in Supplementary Table S2. *Streptomyces* sp. CNT360 and *S. coelicolor* M1152 strains were routinely pre-cultivated in Trypticase Soy Broth (TSB) containing kanamycin and nalidixic acid at 30 °C for three days. A portion (5 ml) of the pre-culture was used to inoculate 500 ml SPM medium at 30°C for seven days. *E. coli* DH10B strains were grown in Luria-Bertani broth (LB) at 37°C. *E. coli* BL21(Gold) strains were cultured in Terrific Broth (TB) at 37°C for general growth and 18°C for protein expression.

### Transformation-associated recombination (TAR) cloning and heterologous expression

The integrative TAR capture vector pCAP01 was linearized by digestion with SpeI and XhoI to yield L-pCAP01. *Streptomyces* sp. CNT360 was cultured in 5 mL TSB and genomic DNA was extracted using standard methods. The *taa* BGC was excised from *Streptomyces* sp. CNT360 genomic DNA by digestion of genomic DNA with PsiI and AclI. Two 1kb homology arms from regions flanking the *taa* BGC in *Streptomyces* sp. CNT360 were amplified by PCR, with primers in each pair containing a short region overlapping one end of L-pCAP01. L-pCAP01 and the two homology arms were then ligated by Gibson Assembly, forming pCAP01_cap. Plasmid pCAP01_cap and the excised *taa* BGC were then co-transformed into *S. cerevisiae* VL6-48 spheroplasts, which were plated on synthetic tryptophan dropout agar. Colonies were screened by PCR to check the left, middle, and right regions (each region about 1 kb) of the *taa* BGC. Subsequently, the positive colonies were cultured in selective liquid media for one day, after which pCAP01 containing the *taa* BGC, pCAP01_*taa*, was extracted. Then, pCAP01_*taa* was first transformed into *E. coli ET12567*, and subsequently conjugated into *S. coelicolor* M1152 by triparental intergenic conjugation facilitated by *E. coli* ET12567/pUB307 to yield the heterologous expression host M1152_*taa*. Cosmids and plasmids used in this study were listed in Supplementary Table S2, and primers were listed in Supplementary Table S3.

### Isolation of compounds 1, 2, and 4

The heterologous expression host M1152_*taa* was pre-cultured from positive conjugant colonies in 100 mL TSB supplemented with nalidixic acid and kanamycin overnight. The pre-culture was diluted 100-fold into twenty 4 L Ultra Yield^®^ flasks with 500 mL SPM in each flask. These 500 mL cultures were incubated with agitation at 30°C for seven days, after which the cultures were centrifuged at 8000 rpm for 10 min at 4°C and the supernatant was collected. The supernatant was adjusted to pH 4.0 using formic acid and was subsequently extracted with equal volume ethyl acetate. The extract was dried, redissolved in ACN, and subjected to semi-preparative RP-HPLC with mobile phase 45% ACN: 55% H_2_O and flow rate 3 mL/min. Fractions containing compounds **1** and **2** based on LCMS analysis were manually combined. The collected fractions were frozen and subsequently lyophilized before being stored at -80°C. Compound **4** was isolated using the same process from heterologous expression host M1152_*taa*Δ*taaH*.

### X-Ray structure determination of 1 coordinated with Cu(II)

X-ray intensity data from a blue needle crystal were collected at 100(2) K using a Bruker D8 QUEST diffractometer equipped with a PHOTON-II area detector and an Incoatec microfocus source (Mo Kα radiation, λ = 0.71073 Å). The raw area detector data frames were reduced, scaled, and corrected for absorption effects using the Bruker APEX3 (Version 2019.11-0), SAINT+ (Version 8.40B) and SADABS programs^47^. The structure was solved with SHELXT^48,49^. Subsequent difference Fourier calculations and full-matrix least-squares refinement against *F*^2^ were performed with SHELXL-2018^48,49^ using OLEX2^50^.

The compound crystallizes in the orthorhombic system. The pattern of systematic absences in the intensity data was uniquely consistent with the space group *P*2_1_2_1_2_1_, which was confirmed by structure solution. The asymmetric unit consists of two crystallographically independent but chemically similar Cu(C_15_H_24_NO_3_S)_2_(H_2_O) complexes, five independent water molecules and three independent ethanol molecules. The major component fractional occupancy refined to 0.721(9). All non-hydrogen atoms were refined with anisotropic displacement parameters. Hydrogen atoms bonded to carbon were in general located in difference Fourier maps before being placed in geometrically idealized positions and included as riding atoms with *d*(C-H) = 1.00 Å and *U*iso(H) = 1.2*U*eq(C) for methine hydrogen atoms, *d*(C-H) = 0.95 Å and *U*iso(H) = 1.2*U*eq(C) for arene hydrogen atoms, *d*(C-H) = 0.99 Å and *U*iso(H) = 1.2*U*eq(C) for methylene hydrogen atoms, and *d*(C-H) = 0.98 Å and *U*iso(H) = 1.5*U*eq(C) for methyl hydrogens. The methyl hydrogens were allowed to rotate as a rigid group to the orientation of maximum observed electron density. Hydroxyl (including ethanolic) and water hydrogen atoms were located and refined isotropically with *d*(O-H) = 0.85(2) Å distance restraints. An additional *d*(H-H) = 1.37(2) Å restraint was used for water hydrogens. The largest residual electron density peak in the final difference map is 0.38 e /Å^-_3_^, located 0.84 Å from S4. The absolute structure (Flack *x*) parameter after the final refinement cycle was 0.009(2), consistent with the establishment of the correct absolute structure and stereochemistry at all chiral centers.

### Derivatization of 2 to generate 3

Compound **2** (9.8 mg, 1.0 equivalent) was dissolved in 1 mL of anhydrous MeOH and TMS-CH_2_N_2_ (50.4 mg, 15 equivalents) in CH_2_Cl_2_ was added dropwise under nitrogen flow (balloon) and stirred at room temperature. The reaction was monitored by Si TLC (DCM:MeOH, 20:1). An additional 125 μL of TMS-CH_2_N_2_ solution was added after 2.5 h, followed by another 250μL of TMS-CH_2_N_2_ solution after 5 h. The reaction stopped in 8 h. After the reaction, the sample was dried under nitrogen air flow. The reaction product mixture was purified using HPLC with a semi-preparative RP-18 column to isolate compound **3**.

### λ-Red-mediated gene deletion to probe biosynthesis

An apramycin resistance (aac(3)IV) cassette was amplified by PCR from plasmid pMXT19 with homology to regions flanking *taaA-D*. The RPC product and pCAP01_*taa* plasmid were then transformed into *E. coli* BW25113/pIJ790 by electroporation. Following λ-Red mediated deletion, the *taaA-D* deletion mutant was grown on LB agar with apramycin for selection, after which positive colonies were screened by PCR. The resulting plasmid pCAP01_*taa*Δ*taaA-D* was conjugated into *S. coelicolor* M1152 to yield the heterologous expression host M1152_*taa*Δ*taaA-D*. The heterologous expression strain was pre-cultured in 5 mL TSB media and subsequently diluted 100-fold into SPM, which was then incubated with agitation for seven days at 30°C. Then, the supernatant was extracted with an equal volume of ethyl acetate and this crude supernatant extract was analyzed via LCMS to identify the putative function of the deleted gene. This process was repeated for deletion of *taa5, taaH, taaI*, and *taaJ* to investigate the functions of each gene in the *taa* BGC.

### Genetic complementation

The *taaH* gene was amplified from *Streptomyces* sp. CNT360 by PCR, along with the overlapped regions of pKY01 plasmid flanking the *taaH* gene. The *taaH* gene was subsequently assembled with the NdeI and HindIII digested vector pKY01 via Gibson Assembly. The resulting plasmid, pKY01_*taaH*, was then conjugated into the *taaH* deletion mutant (M1152_*taa*Δ*taaH*), and the genetically complemented strain (M1152_*taa*Δ*taaH*::*taaH*) was then cultured on MS agar to allow for integration of the complemented *taaH* into the genome.

### Generation of compound 5

Compound **4** (7.0 mg, 1.0 equivalent) and potassium hydrosulfide (3.0 mg, 2.0 equivalents) were dissolved in 50% pyridine/H_2_O (2.0 mL), then stirred for 8 h at 60°C. The aqueous layer was extracted three times with ethyl acetate after quenched with 10% HCl to pH = 1. The combined organic layers were isolated by semi-preparative HPLC (CH_3_CN:H_2_O, 50:50) to afford compound **5** (3.6 mg, yield 51.4%).

### UV-vis spectra of TaaH and cofactors

All UV-vis absorbance spectra were obtained on a Biospectrometer^®^ basic UV-vis spectrophotometer (Eppendorf, Hamburg, DE) at room temperature. All measurements were made in the 300-600 nm range using a μCuvette G1.0 (Eppendorf, Hamburg, DE) with a 1 mm column length using 3 μL of each solution. Solutions containing 150 μM TaaH protein, 1:1 (v/v) 150 μM TaaH with 60 μM FAD, and 1:1 (v/v) 150 μM TaaH and 150 μM NADPH were prepared in storage buffer ((1x PBS + 10% glycerol) immediately prior to each measurement A blank measurement using the storage buffer was taken prior to sample analysis. The resulting spectral data was visualized using the ggplot2 package version 3.5.0 in R version 4.3.3.

### Circular dichroism (CD) spectra of TaaH and TaaH mutants

All CD spectra were obtained on a J-815-150L CD Spectrometer equipped with a Xe lamp (JASCO, Tokyo, JP). Instrument operation and data acquisition were controlled using Spectra Manager™ for Windows 95/NT version 1.55.00 [Build 2] and J-800 Control Driver version 1.27.00 [Build 1]. All measurements were made using 200 uL of solution in a 1 mm quartz cuvette. All enzyme solutions were prepared to 0.25 mg/mL in storage buffer. All spectra were baseline corrected using a blank solution of storage buffer. Peak smoothing for each spectrum was achieved using the Savitsky-Golay filter with a deconvolution width of 15. Original ellipticity measurements were recorded in degrees and were converted to molar ellipticity ([θ]) using the following equation:

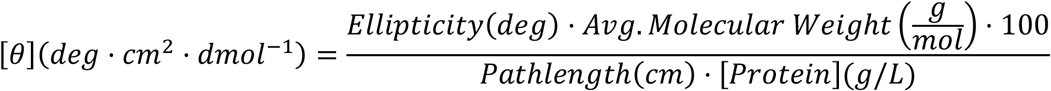

### _34_S-labeled L-cysteine feeding experiment

The native producing strain *Streptomyces* sp. CNT360 was pre-cultured in TSB at 30 °C for three days, then diluted 50-fold into 2 mL SM8 medium (MOPS 20.9 g/L, L-proline 10 g/L, glycerol 20 g/L, NaCl 0.5 g/L, K_2_HPO_4_ 2.1 g/L, EDTA 0.25 g/L, and CaCl_2_ 0.029 g/L, pH 6.5) and cultured at 30°C. On the second day of cultivation in SM8, ^34^S-labeled L-cysteine (0.1 mg/mL) was added directly to the culture. After continuing cultivation for 6 days, the culture was extracted with 2 mL ethyl acetate. The extract was dried, redissolved in ACN, and subjected to HR-ESIMS analysis.

### Plasmids construction for *in vitro* enzymatic assays

To construct plasmids for the expression of TaaI, TaaJ, TaaS1, TaaS2, TaaS3, and TaaS4 with a 6xHis tag at the N-terminus, the genes encoding the corresponding enzymes were PCR amplified from the genomic DNA of *Streptomyces* sp. CNT360 with appropriate overhangs for Gibson assembly. The vector pHis8 was linearized by digestion with NcoI and EcoRI. The linearized vector and the PCR-amplified products were then ligated via Gibson assembly. For the expression of TaaH, the *taaH* gene was amplified from the genomic DNA of *Streptomyces* sp. CNT360 with appropriate overhangs for Gibson assembly. The vector pET28-SUMO was linearized by PCR, and the linearized vector was ligated with the PCR-amplified products via Gibson assembly to construct the plasmid pET28-SUMO-*taaH*. Mutagenesis of the plasmid pET28-SUMO-*taaH* was performed using PCR-based site-directed mutagenesis. Primers incorporating the desired base changes were designed and applied through PCR to amplify the target genes containing the mutations. The mutated PCR products were then used for Gibson assembly with linearized pET28-SUMO, as described above, to generate plasmids containing the mutations. Primers used for PCR were listed in Supplementary Table S3, and the sequence of constructed plasmids was confirmed by DNA sequencing.

### Protein expression and purification

Proteins TaaI, TaaJ, TaaS1, TaaS2, TaaS3, and TaaS4 were produced with an 8×His tag at the N-terminus. The wild-type TaaH and its mutants were produced with a 6×His tag and a small ubiquitin-like modified (SUMO) fusion partner at the N-terminus. *E. coli* BL21 (Gold) cells were transformed with plasmids containing genes encoding corresponding proteins (Supplementary Table S2-S3). A single colony was used to inoculate a 10 mL culture of LB supplemented with 50 mg/L kanamycin. The seed culture was grown for 8 h and used to inoculate 1 L of TB with kanamycin. Cells were grown at 37°C to OD600 ∼0.6-0.8, then cooled to 18°C before IPTG was added to a final concentration of 0.25 mM. The culture was incubated at 18°C for an additional 16 h. Cells were harvested by centrifugation at 5,000×g for 30 min at 4°C. Cell pellets were resuspended in 30 mL of lysis buffer (20 mM Tris, pH 8.0, 300 mM NaCl, 25 mM imidazole, 5% glycerol) and the suspension was sonicated on ice for 20 min to lyse the cells. Cell debris was removed by centrifugation at 15,000 ×g for 60 min at 4°C. The supernatant was loaded onto a 3mL His SpinTrap™ column (GE Healthcare) previously charged with Ni^2+^ and equilibrated in lysis buffer. The column was washed with 10 mL of wash buffer I (20 mM Tris, pH 8.0, 300 mM NaCl, 35 mM imidazole) and 10 mL of wash buffer II (20 mM Tris, pH 8.0, 300 mM NaCl, 55 mM imidazole). The protein was eluted stepwise with elution buffer I (20 mM Tris, pH 8.0, 300 mM NaCl, 250 mM imidazole) and elution buffer II (20 mM Tris, pH 8.0, 300 mM NaCl, 500 mM imidazole). Resulting elution fractions were collected and analyzed by SDS-PAGE. Fractions containing target proteins were combined, then desalted and concentrated using an Amicon Ultra-15 Centrifugal Filter Unit (MWCO, Millipore). The resulting protein samples were stored at −80°C in storage buffer (20 mM Tris, 100 mM NaCl, 10% glycerol).

### *In vitro* enzymatic assays

The enzymatic activity of TaaJ was detected in HEPES (50 mM, pH 7.4) containing **1** (1 mM), CoA (2.5 mM), ATP (2.5 mM), MgCl_2_ (5 mM), TCEP (5 mM), and TaaJ (20 μM) in a total volume of 50 μL. The reaction mixture was incubated at 30 °C for 20 min, then an equal volume of MeOH was added to quench the reaction. To detect the enzymatic activity of TaaI, the reaction mixture was incubated in HEPES (50 mM, pH7.4) containing **1-CoA** (1 mM), Na_2_S_2_O_3_ or L-cysteine (10 mM), ZnCl_2_ (5 mM), TCEP (5 mM), TaaS1 (100 μM), TaaS2 (100 μM), TaaS3 (50 μM), with or without TaaS4 (50 μM), and TaaI (10 μM). The reaction mixtures were incubated at 30 °C for two hours and quenched by ACN. The reaction for TaaH enzymatic activity was performed in HEPES (50 mM, pH7.4) containing compound **5** (100 μM), NADPH (1 mM), FAD (1 mM), TCEP (5 mM), TaaH (10 μM) and incubated at 30 °C for 2 h and quenched by ACN. The enzymatic activities of TaaH mutants were analyzed in the same condition. Boiled WT TaaH and WT TaaH were used as negative and positive control, respectively. The one-pot reactions to synthesize **2** from **1** using a combination of acylsulfenic acid cassette enzymes (TaaJ, TaaI, and TaaH) and primary sulfur donor related enzymes (TaaS1, TaaS2, TaaS3, TaaS4) were performed using **1** (1 mM), Na_2_S_2_O_3_ or L-cysteine (10 mM), CoA (2.5 mM), ATP (2.5 mM), MgCl_2_ (5 mM), NADPH (1 mM), FAD (1 mM), ZnCl_2_ (5 mM), TCEP (5 mM), TaaS1 (100 μM), TaaS2 (100 μM), TaaS3 (50 μM), TaaS4 (50 μM), TaaJ (20 μM), TaaI (10 μM) and TaaH (10 μM) in HEPES (50 mM, pH 7.4) at 30 °C for 2 h and quenched by ACN. All reaction mixtures were filtered after the addition of ACN and analyzed using LC-PDA-ESIMS with gradient elution from 50% to 100% H_2_O/ACN over 10 min, including 0.1% formic acid.

### Kinetics study of TaaH

The kinetic assay was performed in the optimized reaction condition: HEPES (50 mM, pH7.4), NADPH (1 mM), FAD (1 mM), TCEP (5 mM), TaaH (25 nM), and compound **5**. Various concentrations of **5 (**0.001 mM, 0.025 mM, 0.050 mM, 0.075 mM, 0.3 mM, 0.6 mM to 1 mM) were used in the reaction in a total volume of 50 μL. Each reaction was incubated at 30 °C and 50 μL ACN was added to quench the reaction. The reaction mixtures in various time points (0 min, 5 min, 10 min, 20 min, 30 min, 45 min, and 60 min) were analyzed by LCMS as described above. Each kinetic assay was performed in triplicate. The initial rates were calculated by ICEKAT^51,52^. Kinetic enzyme parameters, including K_cat_, K_M_, and V_max_, were calculated using GraphPad Prism (10.1.2).

### TaaH homology modeling and molecular docking

The amino acid sequence of TaaH was analyzed using BLASTP against the Protein Data Bank to identify homologous proteins with known structures. Based on sequence similarity, structure resolution, and the availability of cofactors (FAD, NADP, and oxygen), the crystal structures of flavin-containing monooxygenase from *Pseudomonas stutzeri* NF13 (PDB code: 4usr) and Ancestral Flavin-containing Monooxygenase (FMO) 3-6 (PDB code: 6se3) were selected as templates. Multiple sequence alignment of TaaH with these templates was performed using ClustalW. Homology models of TaaH were generated using Modeller 10.4 by satisfying spatial restraints and followed by structure refinement. Five homology models were generated, and the one with the best DOPE score was retained. Molecular docking of compound **5** with the homology model of TaaH was conducted using GOLD 3.0.1. The docking area was defined by a sphere centered at the ribose moiety (carries the nicotinamide) of NADP, with a radius of 15Å. The genetic algorithm was employed, resulting in ten docking poses. TaaH The best docking pose, based on the docking score, was selected, and its interaction with TaaH was analyzed using Pymol (version 4.6).

### Bioinformatic analyses

Phylogenetic tree analysis was performed using the Neighbor-Joining method. The tree is drawn to scale, with branch lengths in the same units as those of the evolutionary distances used to infer the phylogenetic tree. The evolutionary distances were computed using the Poisson correction method and are in the units of the number of amino acid substitutions per site. All ambiguous positions were removed for each sequence pair (pairwise deletion option). Evolutionary analyses were conducted in MEGA11. The sequence alignment was generated using ClustalW. The sequence similarity network analysis was generated using the online Enzyme Function InitiativeEnzyme Similarity Tool (EFI-EST)^53,54^ and visualized in Cytoscape^55^ with a BLAST *e*-value threshold of 10^−10^. Blast of acylsulfenic acid cassettes was performed using RODEO^56^.

## Data availability

The crystal structure of **1-Cu(II)** has been deposited with the CCDC (https://www.ccdc.cam.ac.uk/) under the CCDC number 2387232. The genome sequence data of *Streptomyces* sp. CNT360 is available in JGI (https://genome.jgi.doe.gov, Project Id: 1016045). Experimental data supporting the conclusions of this study are available within the Article and its Supplementary Information. Source data containing accession numbers and information of proteins used for SSN analysis are provided with this paper.

## Acknowledgements

This work was supported by National Institutes of Health grants 1R35GM150565, R01GM085770, and F32GM129960, as well as a National Science Foundation grant 2239561. We thank Dr. P. Jensen (University of California San Diego, UCSD) and the Fijian government for access to strain *Streptomyces* sp. CNT360 under material transfer agreement 9D00CF1B-436F-4687-A75E-F483ED5B3181. We acknowledge Dr. M. Walla of the University of South Carolina (USC) Mass Spectrometry Facility for assistance in acquiring HRMS and HRMS/MS data, Dr. P. J. Pellechia and Miss T. Johnson from USC NMR Facility, and Dr. B. M. Duggan from UCSD NMR Facility for help with acquiring NMR data, Dr. M. D. Smith from USC X-Ray Diffraction Facility for acquiring X-ray crystallography data, Dr. T. Sawa from Kumamoto University for providing ^34^S-labeled L-cysteine, as well as Dr. K.-S. Ju from The Ohio State University, Dr. Y. Kudo from Tohoku University, Dr. M. S. Cushman from Purdue University, and Dr. Q. Wu from Yale University for helpful discussions.

## Author contributions

D.X., H.Z., B.S.M, and J.L. designed research. D.X., H.Z., W.L., M.D.M., X.L., M.X., C.P., E.A.O., L. H., A.C., T.R., T.A., and C.Y. performed research. All authors analyzed data and discussed results. All authors participated in preparing the manuscript.

